# Extracellular vesicles released by keratinocytes regulate melanosome maturation, melanocyte dendricity and pigment transfer

**DOI:** 10.1101/2023.12.07.570573

**Authors:** Marie-Thérèse Prospéri, Cécile Giordano, Mireia Gomez-Duro, Ilse Hurbain, Anne-Sophie Macé, Graça Raposo, Gisela D’Angelo

## Abstract

Extracellular vesicles (EVs) facilitate the transfer of proteins, lipids and genetic material molecules between cells, and are recognized as an additional mechanism for sustaining intercellular communication. In the epidermis, the communication between melanocytes and keratinocytes is tightly regulated to warrant skin pigmentation. Melanocytes synthetize the melanin pigment in melanosomes that are transported along the dendrites prior to the transfer of melanin pigment to keratinocytes. EVs secreted by keratinocytes modulate pigmentation in melanocytes (Lo Cicero et al., Nat. Comm. 2015). However, whether EVs secreted by keratinocytes contribute to additional processes essential for melanocyte functions remains elusive. Here we show that keratinocyte EVs enhance the ability of melanocytes to generate dendrites, mature melanosomes and their efficient transfer. Further, keratinocyte EVs carrying Rac1 induce important morphological changes, promote dendrite outgrowth, and potentiate melanin transfer to keratinocytes. Hence, in addition to modulate pigmentation, keratinocytes exploit EVs to control melanocyte plasticity and transfer capacity. These data demonstrate that keratinocyte-derived EVs, by regulating melanocyte functions, are major contributors of cutaneous pigmentation and expand our understanding of the mechanism underlying skin pigmentation via a paracrine EV-mediated communication.

**SIGNIFICANCE STATEMENT:** Our work uncovers how keratinocyte-derived EVs control melanocyte physiology and functions. By promoting the growth of melanocyte dendrites, maturation, accumulation and peripheral positioning of pigmented melanosomes within the dendrites, and transfer of melanin to keratinocytes, EVs released by keratinocytes control crucial processes in skin photo protection. Importantly, given that dysregulation of these pathways could underlie pigment disorders, melanoma or skin carcinoma, our results open avenues to exploit keratinocyte EVs as tools for the design of new therapies to enhance the ability of melanocytes to provide skin photoprotection, and thus decrease the incidence pigmentary disorders and skin cancers.

## INTRODUCTION

Skin pigmentation results from the communication between two distinct cell types of the epidermis, namely the pigment producing melanocytes located in the epidermal basal layer and the pigment receiving keratinocytes that are more superficially distributed. Together, they form the epidermal melanin unit in which one melanocyte through its dendrites contacts around 30– 40 keratinocytes (1).This cell-cell physical proximity facilitates the transfer of melanin from melanocytes to the keratinocyte and contribute to skin color and photoprotection against UV solar radiations (2).

Melanin is synthesized in melanosomes, a lysosome-related organelle, which undergoes maturation through four distinct stages (from early non-pigmented stages I/II, to melanized stages III/IV) (3, 4). Maturing pigmented melanosomes undergo microtubule- and F-actin-dependent transport to the tip of melanocyte dendrites where they are captured by actin-related machineries, including the actin-based motor protein Myosin Va (Myo Va), prior to the transfer of melanin pigment to keratinocytes (5–7). Melanosome maturation, dendrite formation and melanosome transfer are stimulated by hormones secreted by keratinocytes including α-melanocyte stimulating hormone (α-MSH), endothelin-1 (ET-1) and nerve growth factors (NGF), as well as ultraviolet radiation. These factors trigger different signaling pathways, such as cAMP/PKA, PKC and MAPK (8–12), that non only increase pigment synthesis, but induces cytoskeletal reorganization, that through the decrease levels of GTP-bound Rho and the concomitant increase Rac1-GTP levels, promote dendrite formation (13–15).

Apart from these soluble factors, keratinocyte release Extracellular Vesicles (EVs) that facilitate communication between keratinocytes and other skin cells, i.e., fibroblasts and melanocytes (16–18). EVs are membrane-bound vesicles that originate from the outward budding of the plasma membrane (ectosomes or large EVs), or from the endosomal network (exosomes or small EVs). EVs carry a variety of proteins, lipids and genetic material [mRNA, microRNAs (miRNAs)] that once transferred to recipient cells, activate different signaling pathways and modulate their fate and behavior (19, 20). In the skin, keratinocyte-derived small EVs, (sEVs) with features of endosome-derived exosomes were shown to modulate melanocyte pigmentation (17, 21, 22). However, the cellular and molecular mechanisms underlying additional functions of sEVs and essential processes that contribute to skin pigmentation, beyond melanin synthesis, remain to be determined.

Here, we have isolated sEVs with features of endosome derived exosomes (17) and adopted the current nomenclature of sEVs according the MISEV guidelines (23). We have investigated sEVs secreted by primary human epidermal keratinocytes (HEKs) to delineate their functional impact on primary human epidermal melanocyte (HEM) activation. We show that HEK sEVs drastically increase the percentage of pigmented melanosomes, promote their accumulation at the dentritic tips, trigger an increase of MyoVa and enhance their melanin transfer ability. Furthermore, we found that HEK sEVs induce morphological remodeling of the HEM plasma membrane and the formation of numerous dendrites, a process that is mediated by Rac1-containing sEVs. Overall, this study put forward novel functions of HEK sEVs, at least through their Rac1 content, in skin pigmentation by controlling the formation of pigmented and transfer-competent dendrites, crucial for the pigment transfer to HEKs.

## RESULTS

### Keratinocyte sEVs promote pigment transfer from melanocytes to keratinocytes

Skin pigmentation and photoprotection rely on sequential processes that involve synthesis of melanin in HEMs, followed by its transfer to adjacent epidermal HEK (24). Recent reports have shown that sEVs released by HEK enhance melanin synthesis in HEMs (17, 21, 22). To investigate whether these sEVs facilitate melanin pigment transfer from HEMs to HEKs, we first isolated sEVs from HEKs of light (HEK-L) and dark (HEK-D) pigmented skin types and characterized these sEVs according to the MISEV guidelines (23) (Fig. S1 and Methods). Immunoblotting of both sEV fractions was positive for classical sEVs markers (Alix, CD63, TSG101, Syntenin), some of which were enriched compared to the corresponding whole-cell lysates (L) and tubulin protein levels (Fig. S1A). HEK-sEVs fractions were devoid of intracellular contaminants as evidenced by the absence of the Calnexin, an endoplasmic reticulum resident protein (Fig. S1A). Nanoparticle tracking analysis (NTA) revealed that both sEV fractions displayed a similar concentration of particles (Fig.S1B). Further visualization of the sEV fractions using transmission electron microscopy (TEM) with negative staining showed that both fractions contained a mixture of cup-shaped sEVs, with various electron densities (Fig. S1C), however, with similar diameter (sEV-L: 86.6 ± 15.8 nm, sEV-D: 84.2 ± 18.9 nm; Fig. S1D). By immuno EM (IEM), we observed that both sEVs fractions were similarly labeled for CD9, CD63, or CD81 sEV-components (Fig. S1E). We further separated the sEVs by velocity iodixanol (OptiPrep) density gradient to control for the presence of potential contaminants co-isolated with sEVs during the ultracentrifugation (Fig. S1F). We found that CD63 was present in two to three consecutive fractions (1.082-1.095 g/ml) whose concentrations range between 7 × 10^9^ and 8.7 × 10^9^ particles/ml for sEV-L or sEV-D respectively (Fig. S1F). These data show that independently of their skin phototypes, HEKs secrete a homogenous population of sEVs in similar quantity and quality.

Next, we investigated the functional effects of HEK sEVs on the transfer of melanin pigment from HEMs to HEKs. HEM-HEK were co-cultured and we analyzed the transferred pigments by immunofluorescence microscopy (IFM). HEMs previously incubated for 48h or not with HEK-sEVs were co-cultured with HEKs for 48h, chemically fixed and co-labelled with anti-EGFR antibody (staining HEKs; green) and HMB45 antibody (staining PMEL fibrils onto which melanin deposits; red) (25). Note that melanized HEMs are strongly labeled for HEM45 as compared to transferred melanin pigment in HEKs (Fig. 1A). When HEKs were co-cultured with sEVs-incubated HEMs (Fig. 1A, right panel), the proportion of HEKs positive for at least one HMB45-positive structure was substantially increased nearly 2-fold compared to control conditions (Ctrl: 30 ±1.5%; sEVs: 60 ± 3.2%, Fig. 1B). This points towards the ability of sEVs to enhance the transfer of melanin pigment from HEMs to HEKs. To reinforce this observation, and as more than one HBM45-positive structure can be transferred to HEKs, we measured the surface area covered by HMB45-positive structures and normalized it to the cell surface area (see Methods). The significant increase of the area covered by HMB45-positive structures per HEKs co-cultured with sEV-incubated HEMs indicates that a higher number of HMB45-positive structures were transferred to each HEK compared to control (Fig. 1C). Taken together, these data show that sEV-treated HEMs exhibit enhanced melanin transfer capacity, allowing a greater number of neighboring HEKs to obtain more melanin pigment, and further indicate that sEVs released by HEKs are facilitators of melanin transfer.

**Figure 1.**
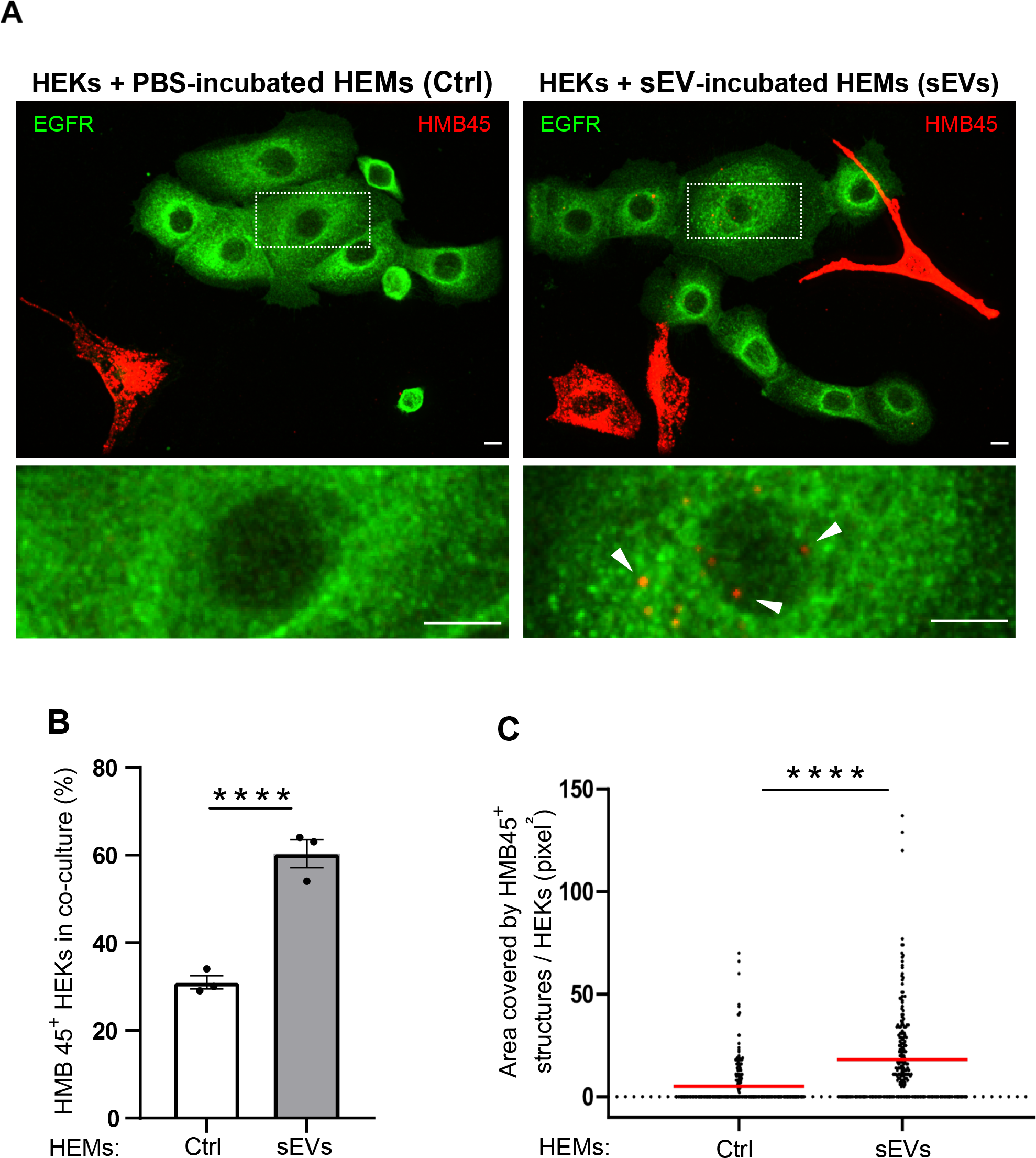
Keratinocyte derived sEVs contribute to the transfer of melanin pigment from melanocytes to keratinocytes. *(A-C)* HEMs treated with HEK sEVs (sEVs) or PBS (Ctrl) for 48h were co-cultured with HEKs for 2 days. *(A)* IFM images of co-cultures immunolabelled for HMB45 (premelanosome protein, PMEL, red) to label melanin pigment, and EGFR (green) to label HEKs. HMB45 staining in HEKs corresponds to transferred melanin pigment. Boxed regions at higher magnification depict the transferred pigment (arrowheads). Bars: 10 μm. *(B)* Quantification of the percentage of HEKs with HMB45-positive structures in each condition. Values are the mean ± SEM of three independent experiments. **** *P* < 0.00001. (Ctrl, n = 238; sEVs, n = 252). *(C)* Quantification of the area covered by HMB45-positive structures per HEK normalized to the total HEK surface area in each condition. Data shown are the results of three independent experiments with the median indicated in red **** *P* < 0.0001. (Ctrl, n = 238; sEVs, n = 252).

### Keratinocyte sEVs promote melanosome maturation, peripheral accumulation and an increase of Myo Va at the tip of melanocyte dendrites

Given that melanosome maturation, accumulation and capture at the dendritic tips are critical steps preceding melanin pigment transfer to neighboring HEK, we set out to examine whether these processes are also affected by HEK sEVs. HEMs were incubated for 48h with PBS (Ctrl) or HEK sEVs (sEVs) from donors of the same phenotype and maturation of melanosomes was analyzed by TEM. Melanosome stages, from unpigmented (stage I and II) to pigmented stages (stage III and IV), were defined based on their morphology and melanin content (Fig. S2A). We observed that mature melanosomes (late stages) were distributed throughout the dendrites of sEV-treated HEMs (Fig. S2A, B right panel), as compared to control, in which mostly unpigmented melanosomes (early stages) were seen (Fig. S2A, B left panel). Remarkably, sEV-incubation promoted a significant decrease in the percentage of unpigmented melanosomes (58% ± 4.6%) concomitant with a 3.2-fold increase in the proportion of pigmented melanosomes (42% ± 4.6%) compared to control (Fig. 2A, B). In contrast, in the absence of sEVs in the medium, the vast majority (90% ± 4%) consisted of unpigmented melanosomes (early stages) while only a small percentage (10% ± 4%) were pigmented (Fig. 2A, B). When we examined in particular the distribution of melanosomes at the dendritic tips of control HEMs, we found only a small proportion of mature melanosomes (10% ± 5.1%). In contrast, exposure of HEMs to HEK sEVs for 48h, triggered a striking increased (3.8-fold) in the percentage of pigmented melanosomes (48% ± 6%) at the dendritic tip, in peripheral position and in close proximity to the plasma membrane compared to unexposed cells (Fig. 2A upper panel, 2C). Interestingly, the overall increase of pigmented melanosomes in sEV-incubated HEMs was similar to the increase of peripheral pigmented melanosomes at the tip of dendrites (Fig. 2B, C).

**Figure 2.**
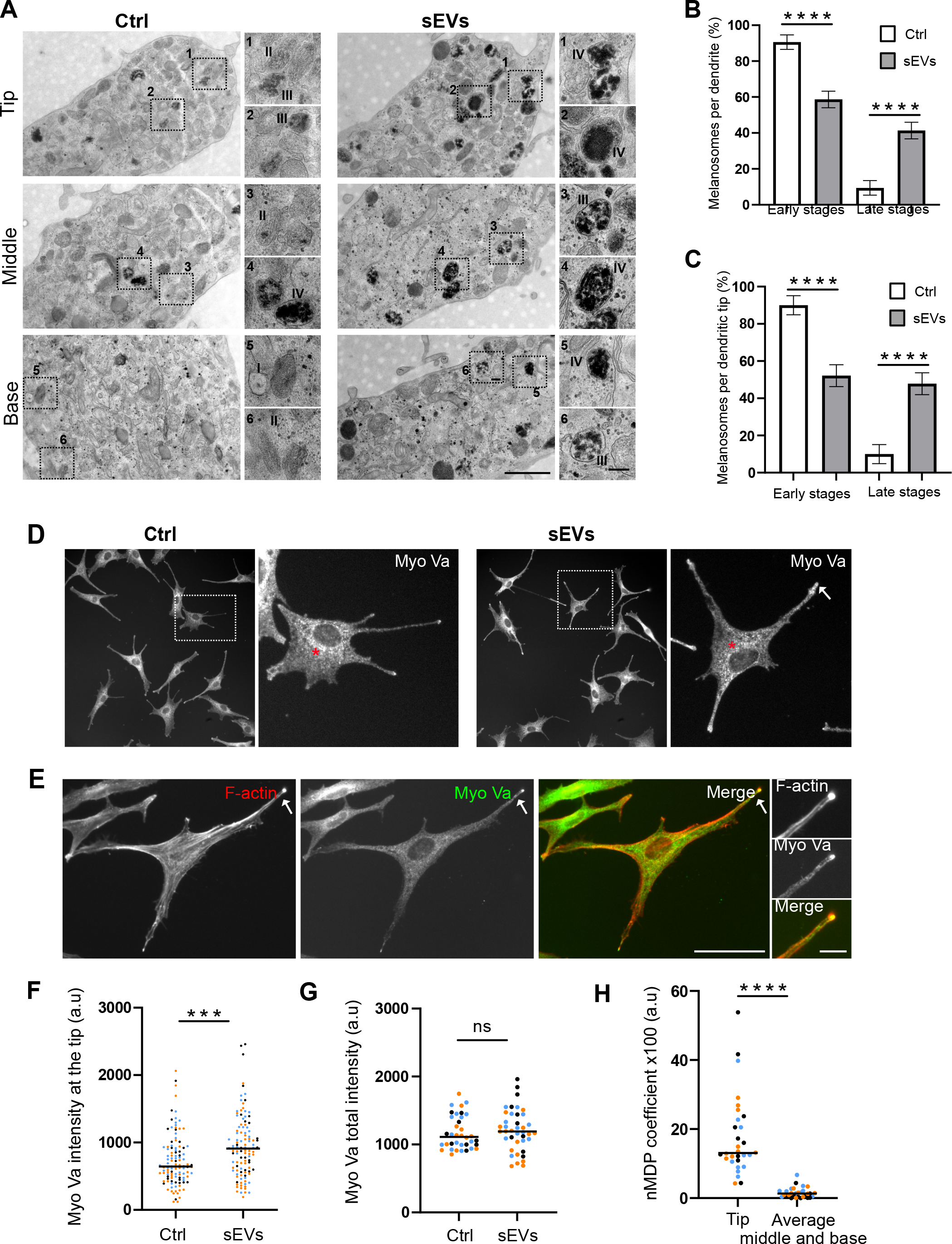
Keratinocyte sEVs induce melanosomes maturation, peripheral accumulation and increased Myo Va at the tip of melanocyte dendrites. *(A)* Representative conventional TEM micrographs of the tip, middle and base of HEM dendrites incubated for 48h with PBS (Ctrl) or HEK sEVs (sEVs). Insets show magnification of the boxed areas. I to IV depict different stages of maturation of melanosomes. Bars: 1μm; insets: 200 nm. (See also Fig. S2A, B). *(B, C)*. Percentage of melanosomes at early and late stages of maturation *(B)* within the dendrites and (*C*) at the dendritic tip of HEM treated as in *(A). (B)* Ctrl: n = 291 melanosomes, 5 dendrites; sEVs: n = 262 melanosomes, 6 dendrites. *(C)* Ctrl: n = 117 melanosomes, 5 dendrites; sEVs: n = 113 melanosomes, 6 dendrites. Values are the mean ± SEM (****P < 0.00001). *(D)* IFM images of HEMs treated as in *(A)*, immunolabelled for Myo Va (white). Red asterisks point to the punctate Myo Va staining throughout the cytoplasm. Bars: 50 μm; insets: 10 μm. *(E)* IFM image of HEMs treated with sEVs for 48h, stained for F-actin (phalloidin, red) and immunolabelled for Myo Va (green). Note the colocalization of Myo Va and F-actin at the dendritic tips (white arrows; inset). Bar: 50 μm; inset: 10 μm. *(F)* Scatter dot plot of Myo Va intensity at the dendritic tip of HEMs incubated as in *(A)*. Only HEMs with three dendrites were considered. Data shown are the results of three independent experiments (a colour per experiment), with the median indicated (n = 112 dendrites). *** *P* = 0.0002. (*G*) Scatter dot plot of Myo Va intensity in the whole HEMs (Myo Va total intensity) incubated as in *(A)*. The same HEMs as in *(F)* were considered. Data shown are the results of three independent experiments (a colour per experiment), with the median indicated (Ctrl, n = 36 cells; sEVs, n=40 cells). n.s. *P* = 0.4735. *(H)* Scatter dot plot of the normalized Mean Deviation Product coefficient n(MDP) for colocalization of Myo Va with F-actin in HEMs incubated for 48h with HEK sEVs which exhibited significant accumulation of MyoVa at dendritic tips compared to control (see D). Comparison of nMDP values at the tip and average nMDP values at the middle and base of the dendrites. Data shown are the results of three independent experiments (a colour per experiment), with the median indicated (n = 30 dendrites). **** *P* < 0.0001.

Myo Va binds to F-actin filaments and facilitate thereby the positioning and anchoring of pigmented melanosomes at the dendritic tip previous to the transfer of melanin pigment to HEKs (5–7). We speculated that the accumulation of pigmented melanosomes at the dendritic tips of sEVs-treated HEMs might be linked to an increase of Myo Va/F actin colocalization. To test this, HEMs were first immunolabeled with Myo Va and then analyzed by IFM. We found that in both conditions, Myo Va staining was punctate throughout the cytoplasm (Fig 2D, red asterisk). Interestingly, Myo Va accumulated at the dendritic tip when HEMs were treated with HEK sEVs (Fig. 2D, white arrow). However, when HEMs were incubated with HEK sEVs, quantitative analysis of the signal distribution revealed that sEV-exposure specifically promoted a substantial increase in Myo Va intensity at the dendritic tip but did not alter the overall Myo Va intensity in sEV-treated HEMs compared to control (Fig. 2D white arrow, F, G). This shows that sEVs support the distribution of Myo Va at the tip of HEM dendrites.

Next the double staining for Myo Va and F-actin showed that at the uppermost tip of sEV-treated HEM dendrites MyoVa massively colocalizes with subcortical actin (Fig. 2E, white arrow). This was further confirmed by the comparison of normalized Mean Deviation Product (nMDP) values at the tip of dendrites, with the mean nMDP values at the middle and the base of the same dendrite (Fig. 2H, Fig. S2D). All together these findings show that HEK sEVs drastically increase melanosome maturation, their accumulation, the build up of Myo Va at the distal end of the dendrites, and Myo Va/subcortical actin colocalization. Thus, it strongly suggests that HEK sEVs facilitate the peripheral positioning of pigmented melanosomes likely through specific tethering at the dendritic tips.

### HEK sEVs promote dendrite outgrowth in melanocytes

Given that HEK sEVs promote melanosome maturation, accumulation, and positioning of melanosomes as well as their transfer to keratinocytes, we next investigated whether sEVs could also promote *per se* dendrite formation. In HEMs, the increase of cAMP levels, by promoting cell shape alterations, supports dendrite outgrowth (13, 14, 26). To functionally test the effect of sEVs on the generation of dendrites, we first performed a set of experiments in which we stimulated HEMs with forskolin (FSK) a pharmacological factor that up-regulates the cAMP pathway and known to change HEM morphology (13, 14, 26) followed by IFM analysis using α-tubulin and F-actin immunostaining. To evaluate HEM dendricity, we focused exclusively on HEMs with defined extensions emanating from the cell soma (Fig. 3A, white arrows), measured their diameter and considered the diameter of dendrites to be less than or equal to 2 μm, while that of protrusions was larger. We observed that the majority of non-stimulated HEMs (DMSO-treated) display the characteristic bipolar morphology with thick protrusions (Fig. 3A, left panel, red arrows). In contrast, FSK stimulation triggered morphological changes (Fig. 3A, right panel, blue arrow), decreased the number of HEMs with protrusions (61% ± 6% vs 78% ± 8% for DMSO-treated) while increasing the number of dendritic HEMs (88% ± 4% vs 63% ± 9% for DMSO-treated cells) (Fig. 3B, C, two first bars). Moreover, the percentage of FSK-treated HEMs with 2 dendrites was 2.5-fold the percentage for DMSO-treated cells (Fig. 1D tenth and ninth bar). When HEM were incubated with HEK sEVs, HEMs displayed a stellate morphology with dendrites (Fig. 3A, bottom, right panel, blue arrow), whereas most of the control HEMs (PBS-treated) (75% ± 4%) exhibited a spindle shape with protrusions (Fig. 3A, bottom, left panel, red arrow; B third bar), and 63% ± 7% (Fig. 3C third bar), had one or two dendrites (42% ± 3% and 21% ± 4% respectively) (Fig. D seventh and eleventh bar). Importantly, incubation with sEVs promoted a marked increase of the number of dendritic HEMs (about 91% ± 5%) (Fig. 3C, last bar) while simultaneously reducing the number of cells with protrusions (about 57 % ± 2%) (Fig 3B, last bar). sEV-incubation also stimulated de novo dendrite formation with a higher percentage of cells having 2 and 3-4 dendrites (1.7-fold and 6-fold respectively higher that in control cells; Fig. 3D, twelfth and last bar). To further quantify the observed morphological changes, we used a second approach to measure the Aspect Ratio (AR) (methods). Reinforcing our findings, the effect of sEVs on HEM dendricity was similar to the effect of FSK and correlated with a significant reduction of the AR ratio, and thereby a higher cell spreading (Fig. 1E). Altogether, these data show that the sole incubation with sEVs is sufficient to alter HEM morphology and support dendrite outgrowth to a comparable level to that of FSK. We also noted that the ability of HEK sEVs to promote such a dendricity was an intrinsic feature of HEK and unrelated to their phototypes, as dendrite outgrowth was also observed in HEMs of dark phototype incubated with sEVs from HEKs from donor of the same phototype (Fig. S2C). Taken together, these findings underscore the correlation between dendrite formation and morphological changes, and indicate that HEK sEVs, independently of HEK skin phototype, bear components that regulate melanocyte function. More importantly, these results demonstrate for the first time that HEK sEVs are potent stimulators of HEM dendricity and strongly suggest a possible effect of HEK-sEVs on HEM cytoskeleton organization.

**Figure 3.**
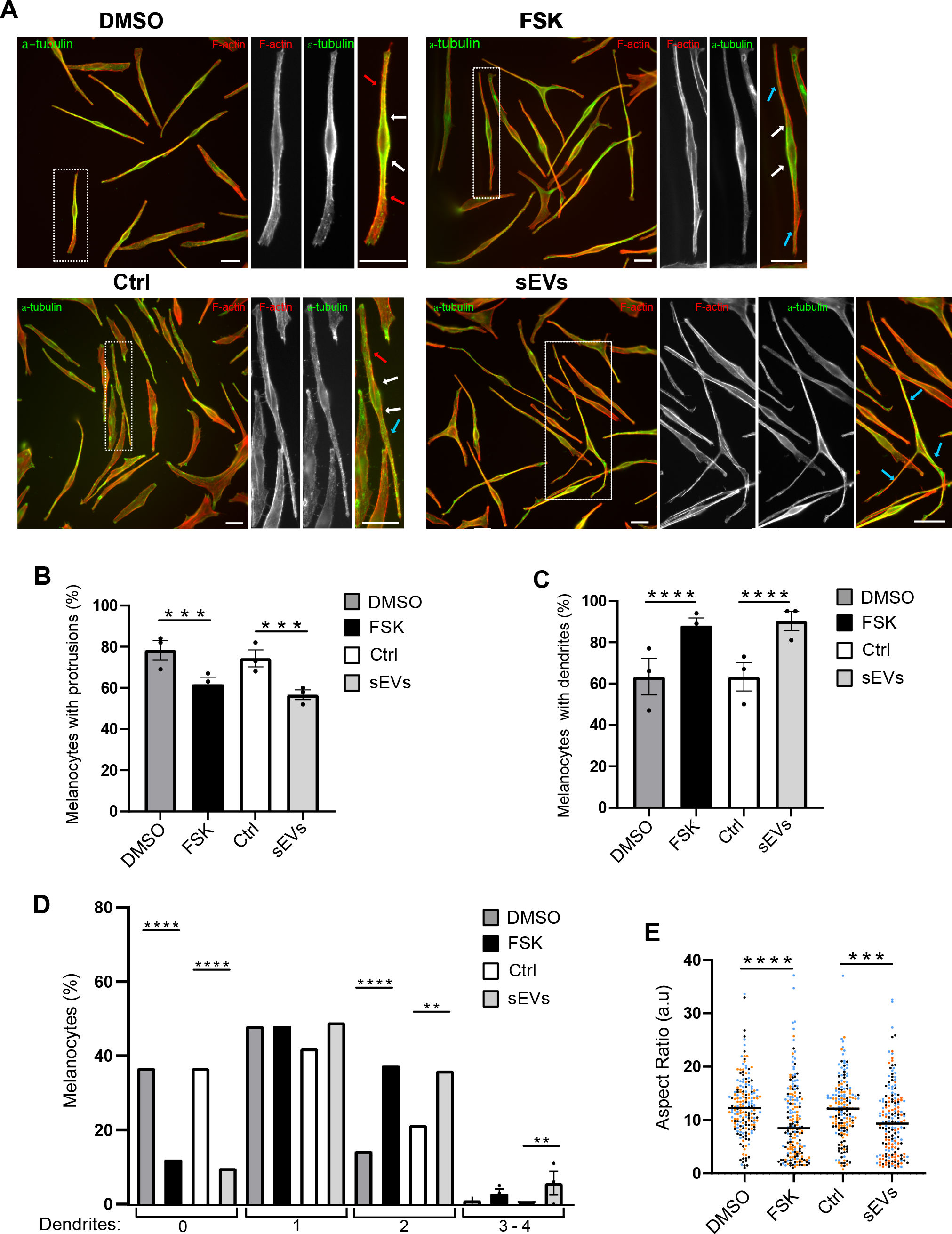
Keratinocyte sEVs alter melanocyte morphology and promote dendrite outgrowth. *(A)* IFM images of HEMs treated for 24h with DMSO or FSK, or for 48h with PBS (Ctrl) or HEK sEVs (sEVs), immunolabelled for α-tubulin (green) and stained for F-actin (phalloidin, red). The boxed regions mark the zoomed area. Bars: 25 μm. Extensions emanating from the cell soma, white arrows; protrusions, red arrows; dendrites, blue arrows. *(B)* Percentage of HEMs [treated as in *(A)*] harbouring protrusions (DMSO, n = 186; FSK, n = 187; Ctrl and sEVs, n = 177). Values are the mean ± SEM of three independent experiments. Only significant *P* values are indicated. *** *P* = 5 × 10^−4^. *(C, D)* Percentage of HEMs [treated as in *(A)*] showing dendrites *(C)*, or 0, 1, 2 or 3 dendrites *(D)* (DMSO, n = 186; FSK, n = 188; Ctrl, n = 178; sEVs, n = 183). Values are the mean ± SEM of three independent experiments. Only significant *P* values are indicated. *(C)* **** *P* = 5.5 × 10^−5^. *(D)* 0 dendrites **** *P* = 5.5 × 10^−5^. 2 dendrites, DMSO and FSK, **** *P* =5.3 × 10^−5^; Ctrl and sEVs, ** *P* = 0.008. 3 dendrites, ** *P* = 0.0032. *P* values are calculated by comparing the number of dendrites in each category to the sum of the number of dendrites in the other categories. *(E)* Scatter dot plot of the AR of HEMs. Data shown are the result of three independent experiment (a colour for each experiment) with the median indicated (DMSO, n = 186; FSK, n = 188; Ctrl, n = 177; sEVs, n = 183). Only significant *P* values are indicated. DMSO and FSK, **** *P* < 0.0001; Ctrl and sEVs *** *P* = 0.0004.

### Melanocyte dendricity is mediated by Rac1-containing exosomes

The initial process of dendrite formation in human melanocytes requires the cytoskeleton reorganization (Evans et al. 2014) mediated by the cAMP-dependent activation of the small GTP binding protein Rac1 (11, 13, 14). As Rac1 protein is expressed (17) and sequestered in HEKs sEVs (Fig. S3A lane 1), we wondered whether Rac1 expression level in HEK sEVs contribute to promote HEM dendricity. First, we evaluated the consequences of Rac1 depletion on HEK morphology and on the quality and quantity of sEVs they secreted using same experimental approaches as described above (Fig. S1). HEKs were transfected with siRNAs targeting Rac1 (siRac1) or control siRNA (siCtrl). We found that Rac1 silencing significantly reduced Rac1 protein and Rac1 mRNA levels as compared with control in cell lysates and sEVs (Fig. S3A, B). Rac1-depleted HEKs harbored a similar morphology as siCtrl-treated HEKs (Fig. S4A), suggesting that depletion of Rac1 did not significantly affect the cytoskeleton and membrane dynamics capacity of HEKs [as shown by IFM (F-actin) and EM; Fig. S4 A, B]. In addition, sEVs released by Rac1-depleted HEKs, were equivalent in number and size compared to control (Fig. S3C, D). In agreement, the number and diameter of MVBs relative to control were similar (Fig. S4C, D). Reinforcing these findings, qualitative, quantitative and IEM analysis revealed the presence of CD9, CD63 and CD81 in siRac1-sEVS to a comparable extent to siCtrl-sEVs (Fig. S3E). We conclude that depletion of Rac1 in HEKs did not impact the general features of sEV such as biogenesis, morphology, quantity, secretion.

Next, we investigated whether the capacity of HEK sEVs to facilitate HEM dendricity could rely on their Rac1 content. HEMs were incubated with PBS (Ctrl), or sEVs released by siCtrl-or siRac1-transfected HEKs. As expected from data presented in Fig. 3, HEMs incubated for 48h with siCtrl-sEVs caused an increase in the average percentage of HEMs harboring dendrites compared to PBS treated HEMs (Ctrl) (71% ± 4% vs 50% ± 3% respectively) (Fig. 4A, B). Especially, the percentage of HEMs with no dendrites decreased while those with 2 or 3 dendrites increased (Fig. 4C), as also reflected by the marked reduction of the aspect ratio of HEMs in siCtrl-sEV relative to PBS-treated cells (Fig. 4D). In contrast, HEMs treated with siRac1-sEVs lost their ability to promote HEM dendricity. Remarkably, all the analyzed parameters returned to control levels (Fig. 4 A-D). The results show that HEK-sEVs with reduced Rac1 content fail to promote HEM plasticity and dendrite formation. Taken together, these data substantiate that the molecular content of HEK sEVs, and in particular Rac1, is integrated by HEMs to control dendrite formation, a hallmark of activated HEMs and critical conduits for pigment transfer to HEKs.

**Figure 4.**
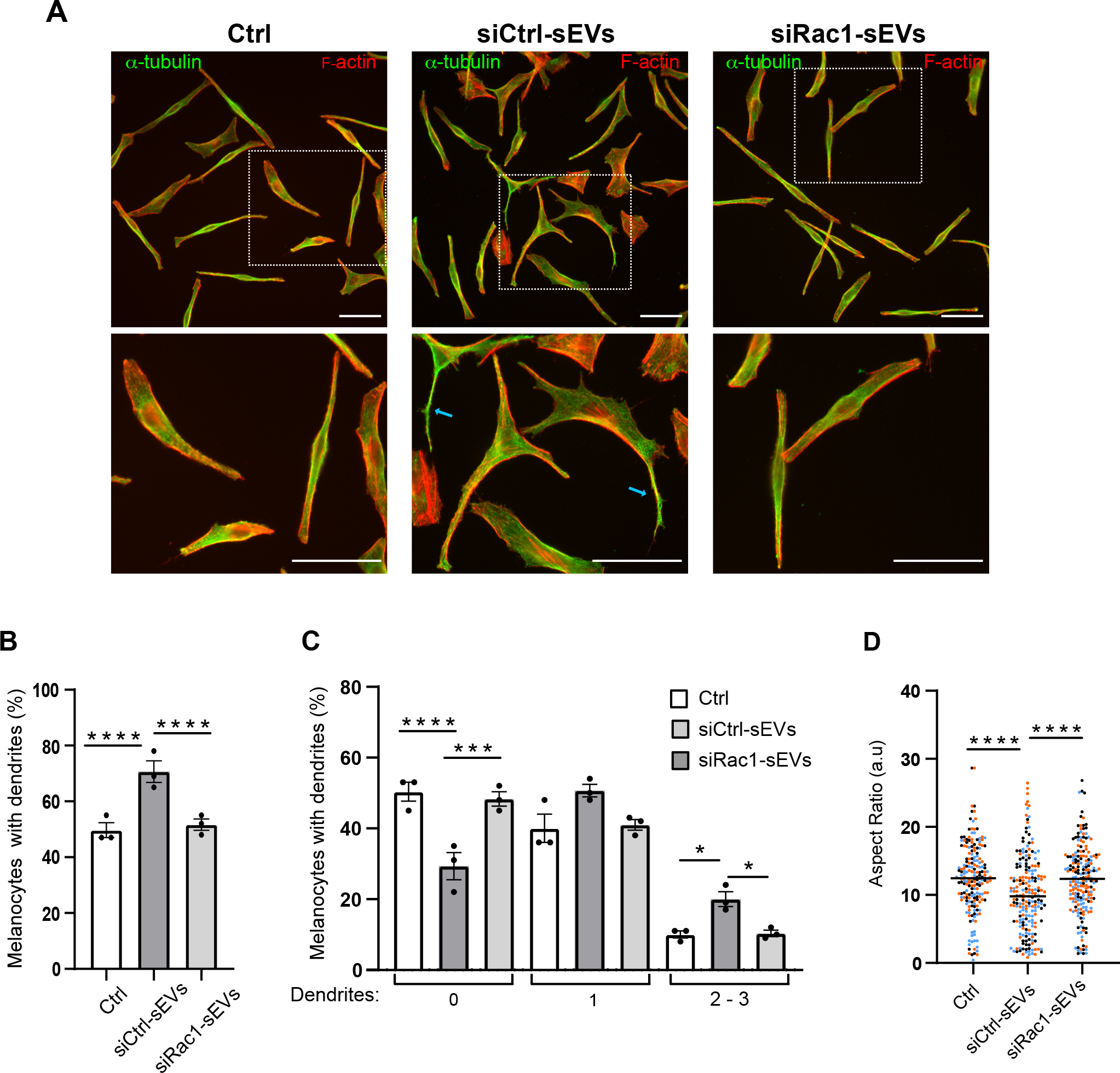
Melanocyte dendricity is mediated by Rac1-containing sEVs. *(A)* IFM images of HEMs incubated with PBS (Ctrl), sEVs from HEK transfected with siCtrl (siCtrl-sEVs), or sEVs from HEK transfected with siRac1 (siRac1-sEVs) immunolabelled for α−tubulin (green) and stained for F-actin (red) with fluorescently labelled phalloidin. The boxed regions mark the zoomed area. Blue arrows depict HEM dendrites. Bars, 50 μm. *(B, C)* Percentage of HEMs [treated as in *(A)*] with dendrites *(B)* or with 0, 1, or 2 - 3 dendrites *(C)* (Ctrl, n = 197; siCtrl-sEVs, n = 206; siRac1-sEVs, n = 204). Values are the mean ± SEM of three independent experiments. Only significant *P* values are indicated. (*B*) Ctrl and siCtrl-sEVs **** *P* = 0; siCtrl-sEVs and siRac1-sEVs **** *P* =0.0001. *(C)* <u>0 dendrite</u>: Ctrl and siCtrl-sEVs **** *P* = 0; siCtrl-sEVs and siRac1-sEVs *** *P* = 0.00045. <u>2 - 3 dendrites</u>: Ctrl and siCtrl-sEVs * *P* = 0.0147; siCtrl-sEVs and siRac1-sEVs * *P* = 0.0191. *P* values are calculated by comparing the number of dendrites in each category to the sum of the number of dendrites of the other categories. *(D)* Scatter dot plot of the AR of HEMs [treated as in *(A)*]. Data shown are the result of three independent experiment (a colour for each experiment) with the median indicated. (Ctrl, n = 197; siCtrl-sEVs, n = 206; siRac1-sEVs, n = 204). Only significant *P* values are indicated. \*\*\*\**P* < 0.0001.

## DISCUSSION

Skin pigmentation, and consequently photoprotection against harmfull UV radiations relies on the production and transfer of melanin from HEMs to HEKs. These processes are tightly regulated by a loop of intercellular communication between both epidermal cells. Several reports showed that soluble factors released by HEKs induce and regulate HEM function, including melanin synthesis and transfer. More recent studies have shown that HEKs release sEVs with features of exosomes carrying selected molecules (e.g. miRNAs), that once internalized by HEMs modulate their pigmentation through an increase in melanin production in a phototype dependent manner (17). In the present study, we demonstrate novel functions of HEK sEVs in stimulating the formation of melanocyte dendrites, maturation of melanosomes and their efficient transfer to HEKs.

We show that HEK sEVs induce changes in melanocyte morphology leading to dendrite formation where mature melanosomes accumulate at their tips, and importantly sEVs promote the transfer of melanin pigment to HEKs. The dendritic process driven by HEK sEVs relies on Rac1 protein which, by modulating actin dynamics and plasma membrane plasticity, endows HEMs with dendrites, special structures for the transfer of melanin pigment to HEKs. These results reinforce and extend previous data showing that HEK sEVs modulate pigmentation via the activation of microphthalmia-associated transcription factor (MITF) (27), and increasing the synthesis and activity of melanosomal enzymes (Tyrp1, Tyr), pigmentation genes such as Rab27 (17).

In the present study we show that HEK sEVs induce significant morphological changes in HEMs and dendrite formation, suggesting that sEVs released by HEK are potent intrinsic activators of dendrite formation. This is firstly due solely to HEK sEVs, as no soluble factors released by HEKs has been added in these experiments, and secondly, a specific feature shared by HEKs independently of their skin phototype. Presumably, HEK sEVs regulate intracellular trafficking machineries and cytoskeletal organization in HEMs through HEK-specific components enclosed and ferried by sEVs. Importantly, treatment of HEMs with HEK sEVs results in changes in cell shape and an increase in dendricity in similar proportions to that of FSK-treatment of HMEs. Since FSK induces an increase in cAMP production in HEMs that is translated into several processes required for pigmentation (melanin synthesis and/or melanosome biogenesis and transport, dendrite formation and/or melanin transfer (this study), the delivery of HEK sEVs to HEMs could represent a powerful alternative intercellular communication pathway for skin pigmentation and photoprotection. Finally, and since high levels cAMP favor the active conformation of Rac1 (GTP-bound Rac1), it is conceivable that part of HEM activation and responses depends on the levels of Rac1/GTP-Rac1 contained in HEK sEVs and subsequently delivered to HEMs. In line with this, sEVs carrying active Rac-1 protein influence recipient cell behavior (28).

Importantly we provide evidence that invalidation of Rac1 in HEKs abolishes sEVs-driven morphological changes and dendrite formation in HEMs. Whether Rac1 sequestered in sEVs is under an active conformation was not investigated due to experimental constraints. However, PKC activation by ET-1 and phorbol esters induces structural changes and HEM dendricity (10, 13) providing a link between PKC and the modulation of Rac1 activity. In addition, PKC might positively regulate the activation of Rac1 through the recruitment and inhibition of Rac1 GTPase-activating proteins (GAPs) (29). Notably, our previous proteomic analysis revealed that HEK sEVs also carry PKC (17). It is thus plausible that the spatiotemporal activation of Rac1 takes place in sEVs prior to their delivery from HEKs. Alternatively activated Rac1 and PKC may be both delivered to HEMs and activate different signaling pathways that would converge and promote HEM dendricity. Therefore, the presence of both Rac1 and PKC in sEVs likely provide a more direct avenue to induce dendrite formation, a pivotal mechanism to expand the contacts with HEMs facilitating thus the transfer of pigment. The effect of HEK sEVs on the remodeling of HEM architecture is certainly a fundamental and a basic feature supporting pigmented melanosome transport and transfer to HEKs. In agreement, sEV-exposure promote melanosome maturation and trigger their accumulation and positioning at the tip of HEM dendrites. This is accompanied by the accumulation of Myo Va and its association with subcortical actin at the dendriti tip, suggesting that sEVs contribute in regulating the actin-based melanosome transport and tethering at the dendritic tip before the transfer to neighboring HEKs. Melanocytes exploit Rab27a/melanophilin/Myo Va for the actin-based transport of mature melanosomes to the actin cytoskeleton in dendritic tips. Myo Va co-localizes with melanosomes (6), and pigment cells overexpressing a constitutively active form of Rac1 causes the accumulation MyoVa at dendritic tips (11). The increase of Myo Va distribution at the tip, its association with subcortical actin and our previous data showing that HEK sEVs increase Rab27a gene expression levels further support the impact of HEK sEVs on HEM pigmentary functions. Together with the presence of Rac1 in sEVs and its role at inducing actin-dependent changes in HEM morphology and melanosome transport along the dendrites, we propose that Rac1 in sEVs is a key component required for the activation of HEM and skin pigmentation.

Melanin transfer to HEKs is a key step in skin pigmentation. The ability of HEK sEVs to promote pigment transfer suggests that sEVs are crucial actors in mediating pigment exchange between these two epidermal cell types. Although the mechanisms of pigment transfer are still debated (30, 31), the generation of dendrites in HEMs is a prerequisite for the establishment of physical contacts with HEKs, while also crucial cellular structures for pigment positioning, enrichment, and transfer, further supporting that sEVs-induced HEM plasticity is pivotal to support these pigmentary functions. In agreement, maintaining the capacity of HEMs to remodel their plasma membrane and to contact HEKs is key for melanin transfer (32). In addition, this process can rely on HEK-secreted factors (soluble factors and EVs) and components, such as miRNAs (32), known to be enclosed in HEK exosomes (17). Therefore, and despite the lack of evidence in 3D-skin tissue system, HEK-derived sEVs may likely cooperate with soluble factors secreted by HEKs to synergize the capacity of HEMs to produce pigment, to provide melanin to HEKs, and to color and photoprotect the skin.

In sum, we propose that sEVs secreted by HEKs and their associated content instruct HEMs so that they integrate and translate these signals into cellular and molecular processes promoting HEM activation. As such, sEVs regulate the expression of melanogenic enzymes and melanosome-associated machineries (17), the remodeling of cell architecture through dendrite formation, the transport and peripheral or pigmented melanosomes to the dendritic tips and importantly, the transfer of melanin pigment to HEKs (this study). Consequently, HEK-derived sEVs are major regulators of the intercellular communication in skin cells and critical actors of the normal skin. Considering that human pigmentation is pivotal to protect the skin against UV solar radiations, and bearing in mind that defects in cutaneous pigmentation lead not only to pigmentary disorders, but can also contribute to several skin cancer types, including melanoma, our study opens up new perspectives for investigating the contribution of the sEVs-mediated HEM functions to such physiological and pathological processes.

## MATERIEL AND METHODS

### Antibodies

The following antibodies were used for immunoblotting (WB), immunofluorescence (IF), or immunoelectron microscopy (IEM): rabbit (rb) anti-β-tubulin (Abcam ab59680; 1:500; WB); rb anti-GAPDH (Sigma G9545, 1:10.000; WB); rb Synthenin (Abcam ab133267;1:1,000; WB); rb Calnexin (Enzo ADI-SPA-860; 1:1,000; WB); rb anti-Alix (Abcam ab186429; 1:1,000; WB); mouse (ms) monoclonal anti-β-actin (Sigma clone AC-74, 1:10,000; WB), ms monoclonal anti-TSG101 (GeneTex, GTX70255; 1:500; WB) or (Santa Cruz, C-2, sc-7964; 1:200 WB); ms anti-Rac-1 (Millipore clone 23A8; 1:1,000; WB); ms anti-human CD81 monoclonal antibody (Millipore CLB579; 1:1,000; WB); ms Anti CD9 (Sigma-Aldrich, clone MM2/57 CLB 162; 1:1,000; WB); ms monoclonal CD63 (Invitrogen, 1:200; WB, IEM); rb anti-MyosinVa (Cell Signaling 3402S; 1:50; IF); sheep anti-EGFR (Fitzgerald 20-ES04, 1:400 IF); ms anti-HMB45 (Abcam ab787; 1:200; IF); ms DM1A α Tubulin (Sigma-Aldrich T6199; 1,200; IF); rb CD9 (Abcam ab236630; 1:80; IEM); rb CD81 (Abcam ab233692; 1:50; IEM); rb anti-mouse Fc fragment (Dako Agilent Z0412; 1:200; IEM). Secondary antibodies coupled to Horseradish peroxidase (HRP)-conjugated goat polyclonal antibodies to rabbit IgG (Abcam ab6721) or to mouse IgG (Abcam ab6789) were used at 1:5,000. Secondary goat anti-rabbit or anti-mouse antibodies conjugated to Alexa Fluor-488, -555 or -647 were from Invitrogen and used at 1:200. Alexa Fluor™ 647 Phalloidin (Thermo Fisher, A22287).

### Cell culture

Normal human epidermal melanocytes (HEMs) and normal human epidermal keratinocytes (HEKs) used in this study were isolated from dark (HEK-D) or light (HEK-L) neonatal foreskin phototypes. HEMs were purchased from PromoCell or Tebu-Bio and HEKs were purchased from CellSystems or Sterlab. HEMs and HEKs were used from passage one to four and maintained in culture in DermaLife Basal Medium supplemented with DermaLife M Life factors (melanocytes-supplemented medium) or with DermaLife K Life factors (keratinocytes supplemented medium) respectively.

To assess HEM dendricity, HEMs (s0 × 10^3^ or 15 × 10^3^ cells) cultured in 0.5 ml of HEM-supplemented medium (followed 5 h later by a PBS rinse) were incubated for 48h in 0.5 ml of HEM-supplemented medium with HEK sEVs (2 × 10^5^ particles/cell or 4 × 10^3^ particles/cell from Optiprep purified sEVs), or with the same volume of phosphate-buffered saline (PBS) as a control (Ctrl), or for the last 24h of the experiment with 30 μM of forskolin (FSK, Sigma-Aldrich F6886), dissolved in dimethylsulfoxide (DMSO, Sigma-Aldrich, CAS 67-68-5), or with 0.6 % DMSO as a control vehicle for FSK. The cells were then processed for microscopy analysis.

To evaluate pigment transfer from HEMs to HEKs, 10 × 10^4^ HEMs cultured in HEMs supplemented medium (followed 5 h later by a PBS rinse) were incubated with HEK sEVs (3 × 10^9^ particles) or PBS (Ctrl) for 48 h in 1 ml HEM-supplemented medium, trypsinized and co-cultured with HEKs at ratio 1:1 in HEM-supplemented medium for 2 days before fixation.

Of note, to be as close as possible to more physiological context, in all experiments HEMs were incubated with sEV derived from HEKs from the same phototype.

### Conditioned Media

The Conditioned Media (CM) was recovered from cultured HEKs. Before recovering the CM, cells were washed once with PBS and cultured subconfluently in fresh HEK-supplemented medium for 48 h in T150 or T75 cell culture flasks.

### siRNA transfection

HEKs (1.25 × 10^5^ cells) were seeded in 100 mm dish and transfected with 40 nM of siRNA using Oligofectamine (Invitrogen) accordingly to manufacturer’s instructions using non-targeting siRNA (si-Ctrl: 5’-AATTCTCCGAACGTGTCACGT-3’), or siRNA targeting the Rac-1 protein (siRac1: 5’-ATGCATTTCCTGGAGAATATA-3’) from Qiagen. Cells were transfected twice; the first shot of transfection 24 h after seeding, followed by a second shot 48 h after the first shot. 4 h after each transfection, the medium was removed and replaced by fresh HEK-supplemented medium (12 ml/dish). CM from each condition was recovered 48h after the second transfection to isolate sEVs.

### Quantitative real-time PCR (qPCR)

Total RNA was extracted from transfected HEKs (i.e., siRNA Ctrl or siRNA targeting Rac1) or HEK sEVs from transfected HEKs (siRNA Ctrl or siRNA targeting Rac1 using the RNA extraction kit (MACHEREY-NAGEL) or Total exosome RNA kit (4478545, ThermoFisher). The cDNA was generated from 0.3 μg of RNA using the SuperScript First-Strand Synthesis System (11904-018, Invitrogen) following manufacturer’s protocols. qPCR was performed using the LightCycler 480 SYBR Green I Master (Roche) on plate-based qPCR amplification and detection instrument LightCycler 480 (Roche). GAPDH was used as an endogenous normalizer. Primers for Rac1: Fw 5’-ATGTCCGTGCAAAGTGGTATC-3’; Rev 5’-CTCGGATCGCTTCGTCAAACA-3’. GAPDH: Fw: 5’-CTGGGCTACACTGAGCACC-3’; Rev 5’-AAGTGGTCGTTGAGGGCAATG-3’. Experiments were performed with at least three biological replicates. The method ΔΔCT was used to obtain the relative expression levels and the ratio between the control and gene of interest was calculated with the formula 2−ΔΔCT.

### Small EVs isolation by differential ultracentrifugation

sEVs were prepared from CM for 48h of 70-90% confluent HEKs. Briefly, CM were centrifuged at 300 g for 10 min at 4°C. Next, supernatant was centrifuged at 2,000 g for 20 min at 4 **°**C to remove debris. Then, the supernatant was centrifuged at 10,000 g for 30 min (4 **°**C) and the sEVs were collected from the supernatant by centrifugation at 100,000 g for 60 min (4 **°**C), in a Ti45 rotor (Beckman). The pellet was resuspended and washed in PBS, pH 7.5 and recentrifuged at 100,000 g for 60 min (4 **°**C), in a Ti70 rotor (Beckman). The resulting pellet was resuspended in PBS and stored at – 80°C.

For the OptiPrep density gradient, a discontinuous iodixanol gradient was prepared, 40, 20, and 10% from a stock solution (Optiprep™ Density Gradient Medium; Sigma-Aldrich; D1556), with 0.25 M sucrose, 10 mM Tris pH 8, and 1 mM EDTA. The 100,000 g pellet was laid at the bottom of the gradient to obtain a 40% fraction and centrifuged at 100,000 g for 18h at 4°C, in a SW41 Ti rotor (Beckman), stopping without brake. Ten fractions of 490 μl each were collected. The fractions were diluted in PBS, centrifuged at 100,000 g in a SW 32 Ti rotor (Beckman) for 60 min at 4°C, resuspended in equal volumes of PBS, and stored at **-**80°C.

### sEV quantification by Nanoparticles Tracking Analysis (NTA)

The isolated sEVs were diluted 1:1,000 in filtered PBS and loaded into the sample chamber. NTA was performed using ZetaView PMX-120 (Particle Metrix) with software version 8.04.02. The instrument settings were at 22°C, sensitivity of 75, and shutter of 75. Measurements were done using two different dilutions, at 11 different positions (two cycles per position) and a frame rate of 30 frames per second.

### Electron Microscopy

#### Transmission Electron Microscopy (TEM) of sEVs

pellets resuspended in PBS were processed as previously described and according to the MISEV guidelines. Briefly, 5 μl of sEVs were deposited on Formvar-carbon-coated electron microscopy grids incubated for 20 min at RT and fixed in 2 % paraformaldehyde (PFA) in Phosphate Buffer (PB 0,2 M, pH 7,4). After 20 min at RT, samples were washed 6 times with H_2_0, and post-fixed in 1 % glutaraldehyde in PBS for 5 min at RT. For negative contrast, the grids were placed on a drop of 4% Uranyl Acetate (UA) and 2% methylcellulose (MC) at 2 % (ratio of 1 UA: 9 MC) for 10 min on ice. The excess was then removed by blotting on Whatman paper and dried for 30 min at RT. All these steps were done in the dark.

#### Immunolabeling electron microscopy (IEM) of sEVs

5 μl of sEVs were loaded on grids and fixed with 2% paraformaldehyde in Phosphate Buffer (PB 0,2 M, pH 7,4) for 20 min at RT. Immunodetection was performed with a mouse monoclonal CD63, a rabbit CD9, or a rabbit CD81 diluted in 1% BSA/PBS for 60 min at RT. Secondary incubation was next performed with a rabbit anti-mouse Fc fragment (Dako Agilent Z0412; 1:200) for CD63 for 20 min, at RT. Grids were incubated for 20 min with Protein A-Gold (PAG) 10 nm diluted in 1%BSA/PBS (1:50) (Cell Microscopy Center, Department of Cell Biology, Utrecht University). A second fixation step with 1% glutaraldehyde in PBS was performed. Grids were contrasted with uranyl acetate and methylcellulose as for TEM. All samples were examined with a Tecnai Spirit electron microscope (FEI, Eindhoven, The Netherlands), and digital acquisitions were made with a numeric 4k CCD camera (Quemesa, Olympus, Münster, Germany).

#### Transmission Electron Microscopy (TEM) of cells

HEKs or HEMs were fixed in 2.5 % (v/v) glutaraldehyde, 2% in 0.1 M cacodylate buffer for 24 h, post-fixed with 1% (w/v) osmium tetroxide supplemented with 1.5% (w/v) potassium ferrocyanide, dehydrated in ethanol and embedded in Epon as previously described (33). Ultrathin EPON sections (60 nm) of cell monolayers were prepared, contrasted with uranyl acetate and lead citrate, and observed at 80 kV transmission electron microscope (Tecnai Spirit G2; Thermo Fisher Scientific) equipped with a 4k CCD camera (Quemesa, Soft Imaging System). Images were analyzed with iTEM software (EMSIS) and statistical analysis was done by GraphPad Prism.

### WB analysis

Cells were lysed on ice in lysis buffer (20 mM Tris pH 7.5, 150 mM NaCl, 1% Triton X-100, 1 mM EDTA, pH 8) with a protease inhibitor cocktail (Roche). Whole-cell lysates or sEVs were incubated in sample buffer with or without DTT (400 mM) (Sigma), boiled for 5 min and fractionated with SDS–PAGE using Nupage (4–12%) Bis-Tris gels (Invitrogen) and transferred to nitrocellulose membranes (Millipore). The membranes were blocked in PBS/Tween 0.1% (PBS/T) with 5% non-fat dried milk, incubated with the indicated primary antibody diluted in the same solution, washed three times in PBS/T and incubated with HRP-conjugated secondary antibody followed by washing in PBS/T. To reveal the membranes, SuperSignal™ West Pico PLUS Chemiluminescent (ThermoFisher) was used.

Images of immunoblots were captured with Chemidoc Touch Biorad 2 or with CL-XPosure Film (ThermoFisher scientific) within the linear range and quantified by densitometry using the “analyze gels” function in ImageJ (http://rsb.ingo.nih.gov/ij/).

### Immunofluorescence microscopy

Cell seeded on glass coverslips were fixed with 2% (v/v) paraformaldehyde in PBS at room temperature for 15 min, and then washed three times in PBS and once in PBS with 50 mM glycine. Dilutions of primary and secondary antibodies were prepared in buffer A: PBS with 0.2% (w/v) BSA and 0.1% (w/v) saponin. The coverslips were incubated with the primary antibodies for 1 h at room temperature after being washed once in buffer A. Coverslips were washed three times in buffer A, followed by incubation with secondary antibodies and phalloidin for 45 minutes if necessary. The final washing steps were performed three times in buffer A, once in PBS and once in water. The coverslips were mounted on glass slides using ProLong™ Gold Antifade Mount with DAPI (Thermo Fisher Scientific).

### Image acquisition

Epifluorescence microscopy was carried out with a Histo-Epifluo Nikon Microscope (Nikon NIe with camera CCD CoolSnap HQ2 and color CMOS sensor DS-Ri2 from Nikon) (Fig. 3A and 4A) and a Leica DM6B microscope equipped with 40x NA 1.3 oil immersion objective and a sCMOS camera (Fig. 2D, 2E, S2D and S4A). Spinning-disc confocal microscopy was carried out with a Yokogawa CSU-X1 on a Nikon Inverted Eclipse TI-E microscope equipped with a 40× NA 1.3 oil immersion objective and a sCMOS Prime 95B (Photometrics) (Fig. 1A). 3D reconstructions were generated by using the software Fiji (https://fiji.sc/).

### Image analysis and quantification

#### Dendrite/protrusion size measurement

The diameter of dendrites or protrusions was measured using the Fiji software’s (34) “straight” tool. Aspect Ratio (AR) was measured using Fiji software’s, “fit ellipse” function which displays the major and minor axis for each cell.

#### Myosin Va and F-actin co-localization

Regions Of Interest (ROI) were manually drawn at the tip of each dendrite, and the same ROI was used to quantify the colocalization values at the tip, middle and base of the same dendrite. Colocalization values were measured using the Colocalization Colormap plugin developed for Fiji (35). This plugin calculates the normalized Mean Deviation Product (nMDP) for each pixel in the image, reflecting the correlation between intensities in the Myo Va (Cy3) and F-actin (Cy5) channels. The nMDP values were normalized by the maximum values.

#### Measurement of Myosin Va intensity

A manually drawn ROI with a diameter of 11.3μm was used to measure in Cy3 images the mean intensity of Myo Va at the tip of each dendrite, as well as the mean background intensity near the same dendrite for all dendrites analyzed. The plotted values correspond to the mean Myo Va intensity at the tip after background subtraction. A manually drawn ROI of the area of each cell was used to mesure in Cy3 images the mean intensity of MyoVa in the whole cell, as well as the mean background intensity near the same cell for all the cells analyzed. The plotted values correspond to the mean total Myo Va intensity in the cell after background subtraction.

#### Quantification of pigment transfer

A Fiji macro was written to perform the analysis on maximum intensity Z projections. The HMB45-positive structures were counted in manually drawn ROI delineating cell contours using ImageJ’s Find Maxima function to identify local maxima. The surface area of these structures was extracted using the Li global threshold method, which minimizes the cross-entropy between the background and the foreground averages, to calculate the proportion of the cell surface area covered by the transferred pigment. HMB45-positive structures at the top of the cells were excluded from quantification.

#### Measurement of Multi-Vesicular bodies (MVB)

Quantification of MVB size and number was determined using the iTEM (Soft Imaging System) software.

### Statistical Analysis

We used GraphPad Prism version 9.0.0 for Windows, GraphPad Software, San Diego, California USA, (www.graphpad.com) for statistical analysis. To compare two conditions (Fig. 1C, 2F, G, H, S3B, D), we performed a non-parametric Mann-Whitney test or umpaired t-test (S1D, S3D, S4C, D). To compare three or more conditions (Fig.3E, 4D), we used a non-parametric Kruska-Wallis test, followed by Dunn’s multiple comparison test. For percentage comparisons (Fig. 1B, 2B, C, 3B-D, 4B,C, S2C), we applied Fisher’s exact test (https://www.socscistatistics.com/tests/chisquare2/default2.aspx), with application of the Benjamini-Hochberg multiple comparison correction (https://www.sdmproject.com/utilities/?show=FDR) when necessary.

## ACKNOWLEDGMENTS

We thank Dr. C. Delevoye, Dr. P. Stahl and Dr. Ph. Vernier for insightful discussions and critical reading of the manuscript; This work was supported by the Centre National de la Recherche Scientifique (CNRS), the Foundation for Medical Research (FRM EQU201903007827), the European Union’s Horizon 2020 research and innovation programme under the Marie Skłodowska-Curie (Grant agreement No 847718)”, and the Cell and Tissue Imaging core facility (PICT IBiSA, Institut Curie), member of the French National Research Infrastructure France-BioImaging (ANR10-INBS-04). C.G. was recipient of a post-doctoral fellowship from the Foundation for Medical Research (FRM SPF201809006932). M.G.D. was supported by PhD fellowship from Institut Curie EuReCa PhD Programme (Grant agreement No 847718).

## Notes

### Competing Interest Statement

The authors have declared no competing interest.

