## Supplemental Figure S1 - S4 for "Extracellular vesicles released by keratinocytes regulate melanosome maturation, melanocyte dendricity and pigment transfer"

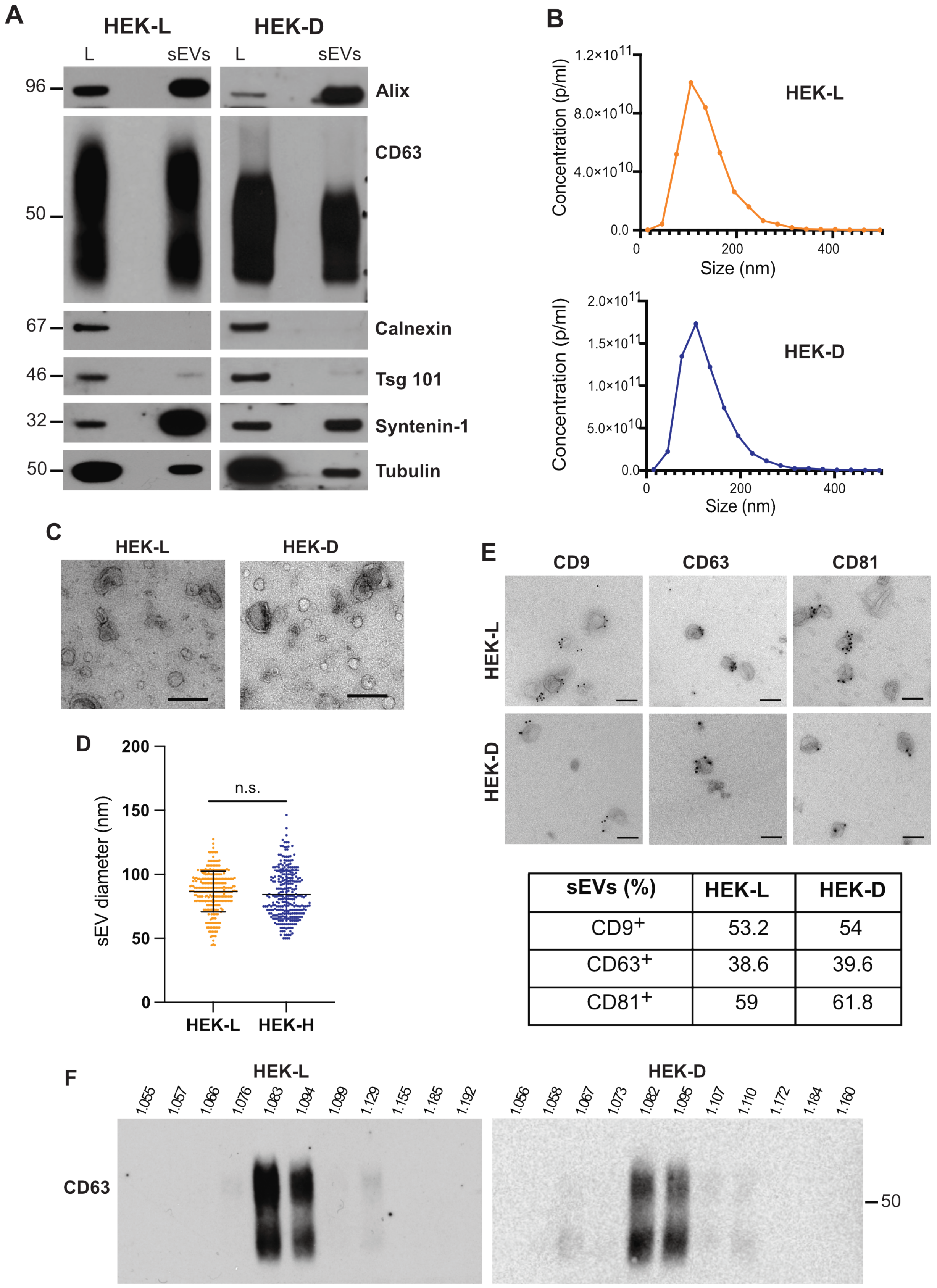


| sEVs (%) | HEK-L | HEK-D |
|---|---|---|
| CD9+ | 53.2 | 54 |
| CD63+ | 38.6 | 39.6 |
| CD81+ | 59 | 61.8 |

**Figure S1: Characterization of sEVs released by HEKs of light (HEK-L) and dark (HEK-D) phototypes.** *(A)* WB analysis of sEVs (Exo; 10 µg) isolated by differential ultracentrifugation from HEK-L or HEK-D phototypes and the corresponding whole-cell lysate (L; 20 µg). The results show the presence of the classical EV marker transmembrane protein CD63, as well as additional exosomal components such as Alix, TSG101 or Syntenin-1, and the lack of the endoplasmic reticulum protein Calnexin, when compared to whole-cell lysates. *(B)* One representative nanoparticle tracking analysis (NTA) of sEVs from HEK-L and HEK-D. *(C)* Representative transmission electron micrograph of sEVs from HEK-L and HEK-D. Bars: 100 nm. *(D)* Scatter dot plot of sEV diameter from HEK-L and HEK-D. Data shown are the results of three independent experiments. Mean ± SEM (n = 286; n.s. $P = 0.0987$). *(E)* Top: Representative images of sEVs from HEK-L and HEK-D immuno-gold labelled for endogenous CD9, CD63 and CD81 (PAG 10 nm). Bars, 100 nm. Bottom: Quantification of the percentage of CD9 positive sEVs (n = 119), CD63 sEVs (n = 183) and CD81 positive sEVs (n = 106). These analyses showed that HEK-L and HEK-D massively release 50-150 nm diameter vesicles partially labelled for CD63 and for two other EV markers, CD9 or CD81. *(F)* sEVs contained in each pellet were further separated by velocity iodixanol gradient and probed with CD63. WB analysis shows that CD63 is present in two to three consecutive fractions (1.082-1.095 g/ml) whose concentrations ranged between $7 \times 10^9$ and $8.7 \times 10^9$ particles/ml for HEK-L or HEK-D respectively.

Figure S2

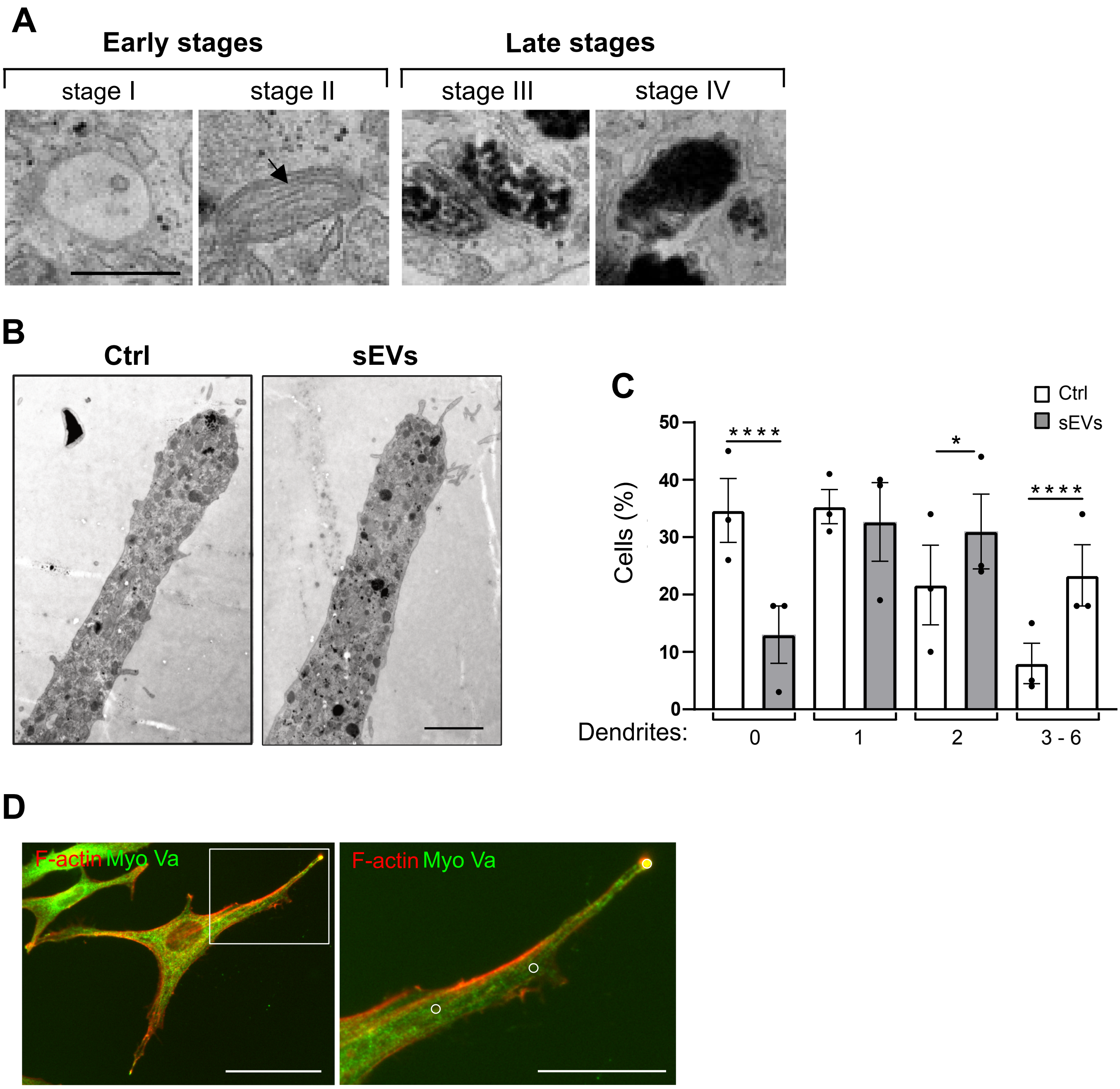
A
Early stages
Late stages
stage I
stage II
stage III
stage IV
B
Ctrl
sEVs
C
Ctrl
sEVs
Cells (%)
50
40
30
20
10
0
****
*
****
Dendrites:
0
1
2
3 - 6
D
F-actin Myo Va
F-actin Myo Va

**Figure S2. Keratinocyte sEVs promote accumulation and peripheral positioning of pigmented melanosomes at the dendritic tips, dendrite formation, and Myo Va/subcortical actin colocalization at the tips.** HEMs were exposed for 48h to HEK sEVs (sEVs) or PBS (Ctrl). *(A)* Conventional TEM micrographs of early unpigmented melanosomes (stages I and II) and late pigmented melanosomes (stages III and IV). At stage I, pre-melanosomes are nonpigmented vacuoles derived from the endosomal system. At stage II, arrow points to internal striations. At stage III, melanin pigment is uniformly deposited. At stage IV, melanosomes are fully mature and pigmented. Bar: 500 nm. *(B)* Representative TEM micrograph of dendrites from control and sEVs-treated HEMs. Note the massive accumulation of pigmented melanosomes at the dendritic tip of sEVs-treated HEM (righ panel). Bar: 2 µm. *(C)* Percentage of HEMs from dark phototype (used to quantify Myo V intensity in Fig. 2 D-G) with 0, 1, 2 or 3 - 6 dendrites (Ctrl, n = 218; sEVs, n = 222). Values are the mean ± SEM of three independent experiments. Only significant *P* values are indicated. 0 dendrites, **** $P$=210$^{-5}$; 2 dendrites, * $P$ = 0.05; 3 - 6 dendrites, **** $P$ = 0. *(D)* Example of the HEM analysed for Myosin Va and F-actin co-localization in Fig. 2E. Note the Regions Of Interest (white circles) drawn at the tip, middle and base of one dendrite (inset). Bars: 50 µm; inset: 25 µm.

### Figure S3

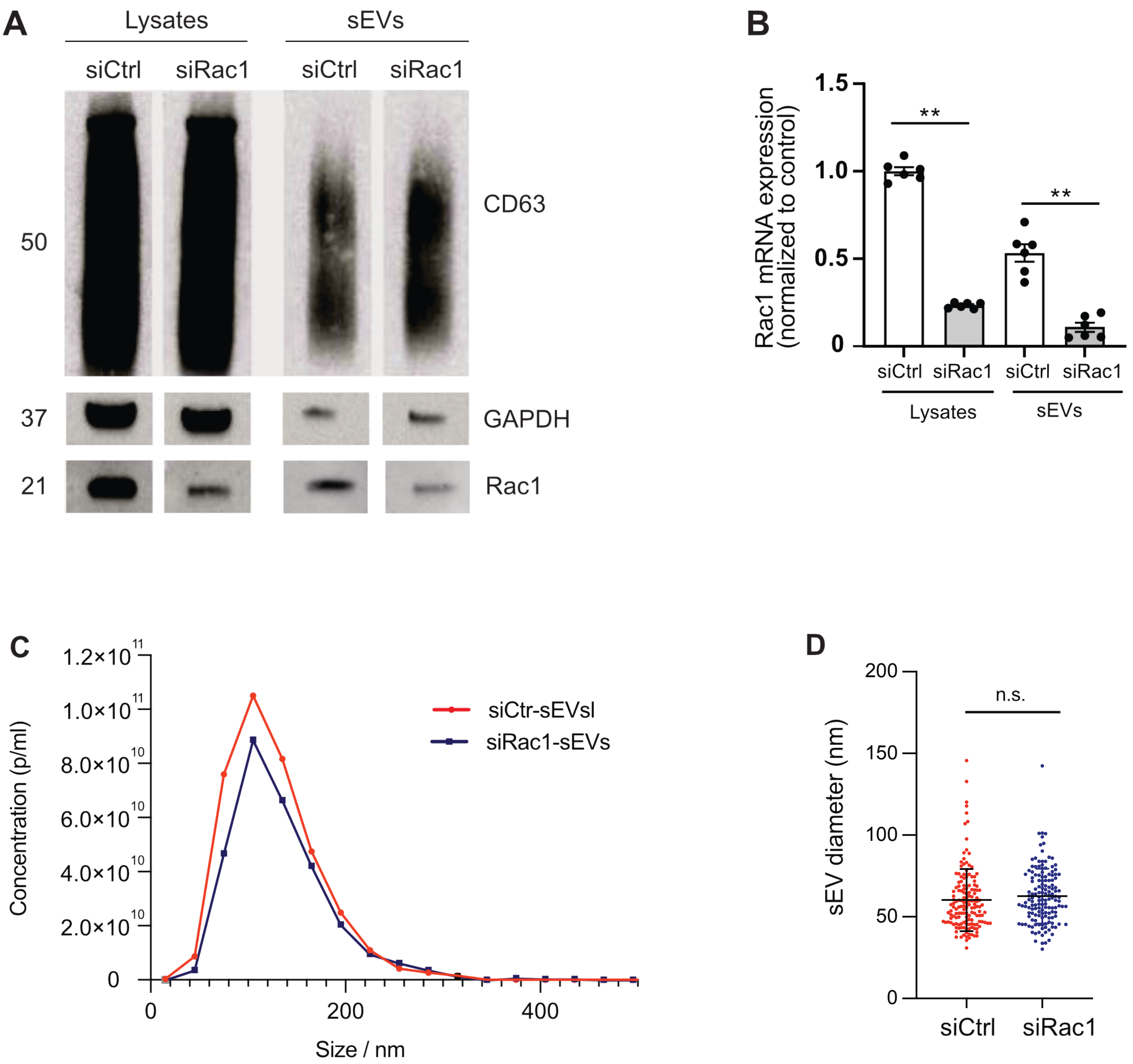


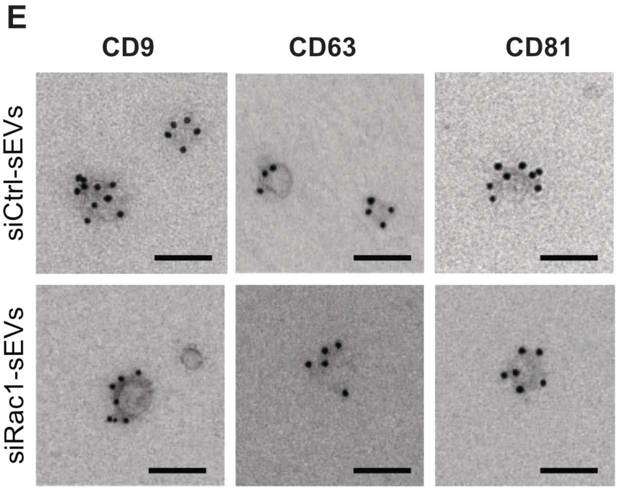


| sEVs (%) | siCtrl | siRac1 |
|---|---|---|
| CD9+ | 49.2 | 39.8 |
| CD63+ | 32.1 | 28.4 |
| CD81+ | 43.6 | 42.2 |

**Figure S3: Characterization of sEVs from siCtrl and siRac1 HEKs.** *(A)* WB analysis of whole-cell lysate (L; 20 µg) and sEVs (sEVs; 10 µg) from siCtrl- and siRac1-transfected HEKs probed with CD63, GAPDH and Rac1 antibodies. Of note, the GAPDH exposure of sEVs is higher than that of cell lysates. *(B)* Quantification of Rac1 mRNA expression normalized to GAPDH in whole-cell lysates and sEVs from siCtrl- or siRac1-transfected HEKs. Values are mean ± SEM of three independent experiments. Only significant *P* values are indicated. ** *P* = 0.0022 for whole-cell lysates and sEVs. *(C)* One representative nanoparticle tracking analysis (NTA) of sEVs from siCtrl- and siRac1-transfected HEKs. *(D)* Scatter dot plot of sEVs diameter. Data shown are the result of three independent experiments. Mean ± SEM (siCtrl n = 151; siRac1 n= 148); n.s not significant. *P* = 0.1215. *(E)* Left: Representative images of sEVs from siCtrl- and siRac1- transfected HEKs immuno-gold labelled for endogenous CD9, CD63 and CD81 (PAG 10 nm) Bars, 100 nm. Right: Quantification of the percentage of CD9 positive sEVs (siCtrl, n = 520; siRac1, n = 585), CD63 positive sEVs (siCtrl n = 327; siRac1, n = 190 and CD81 positive sEVs (siCtrl n = 250; siRac1 n = 279).

### Figure S4

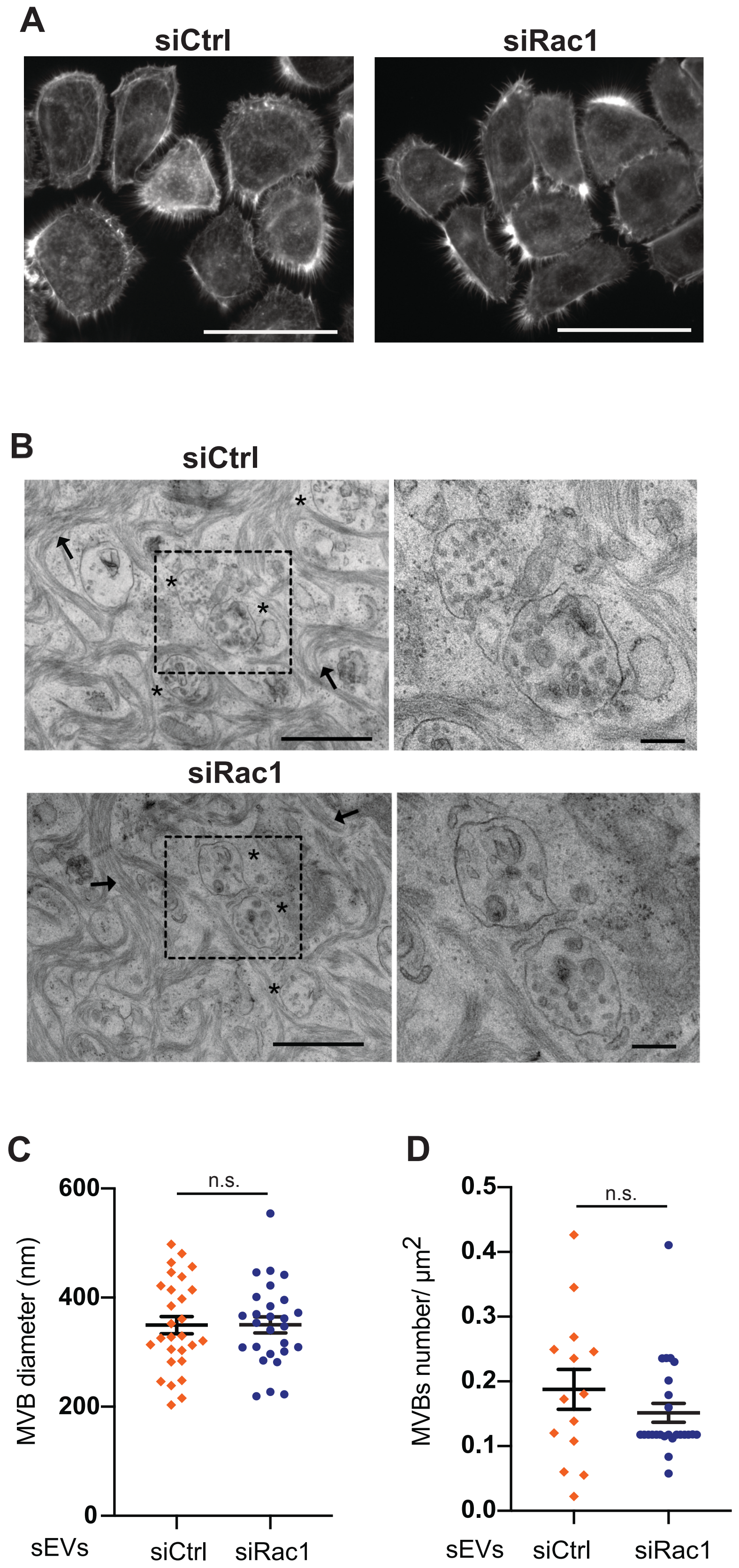

A
siCtrl
siRac1
B
siCtrl
siRac1
C
600
400
200
0
MVB diameter (nm)
n.s.
sEVs
siCtrl
siRac1
D
0.5
0.4
0.3
0.2
0.1
0.0
MVBs number/ μm2
n.s.
sEVs
siCtrl
siRac1

**Figure S4**. **Rac1 silencing does not alter keratinocyte morphology.** HEKs were transfected with siCtrl or siRac1 for 48h and processed for IFM (*A*) or for IEM (*B*). *(A)* HEK were stained for F-actin with fluorescently labelled phalloidin. Bars: 50 µm. *(B)* Conventional TEM micrographs representative of each condition. Black arrows point to keratin bundles. Asterisks to MVBs. Bars: 1 µm. Right panels: magnification of boxed area. Bars, 200 nm. *(C, D)* Scatter dot plot of MVB diameter (C) or MVB number (D) from siCtrl and siRac1 HEK. Data shown are the result of three independent experiments. Mean ± SEM. MVBs were counted on a total surface of 1950 $\mu m^2$. (siCtrl, n = 28; siRac1 n = 27); *(C)* n.s. $P = 0.9820$; *(D)* n.s. $P = 0.2371$.
